# Single-cell transcriptional landscapes of bovine peri-implantation development

**DOI:** 10.1101/2023.06.13.544813

**Authors:** Giovanna Nascimento Scatolin, Hao Ming, Yinjuan Wang, Linkai Zhu, Emilio Gutierrez Castillo, Kenneth Bondioli, Zongliang Jiang

## Abstract

Supporting healthy pregnancy outcomes requires a comprehensive understanding of the cellular hierarchy and underlying molecular mechanisms during peri-implantation development. Here, we present a single-cell transcriptome-wide view of the bovine peri-implantation embryo development at day 12, 14, 16 and 18, when most of the pregnancy failure occurs in cattle. We defined the development and dynamic progression of cellular composition and gene expression of embryonic disc, hypoblast, and trophoblast lineages during bovine peri-implantation development. Notably, the comprehensive transcriptomic mapping of trophoblast development revealed a previously unrecognized primitive trophoblast cell lineage that is responsible for pregnancy maintenance in bovine prior to the time when binucleate cells emerge. We analyzed novel markers for the cell lineage development during bovine early development. We also identified cell-cell communication signaling underling embryonic and extraembryonic cell interaction to ensure proper early development. Collectively, our work provides foundational information to discover essential biological pathways underpinning bovine peri-implantation development and the molecular causes of the early pregnancy failure during this critical period.

**Significance Statement:** Peri-implantation development is essential for successful reproduction in mammalian species, and cattle have a unique process of elongation that proceeds for two weeks prior to implantation and represents a period when many pregnancies fail. Although the bovine embryo elongation has been studied histologically, the essential cellular and molecular factors governing lineage differentiation remain unexplored. This study profiled the transcriptome of single cells in the bovine peri-implantation development throughout day 12, 14, 16, and 18, and identified peri-implantation stage-related features of cell lineages. The candidate regulatory genes, factors, pathways and embryonic and extraembryonic cell interactions were also prioritized to ensure proper embryo elongation in cattle.

## Introduction

Peri-implantation embryo development of ruminant species such as cattle is poorly understood and not closely paralleled in model organisms, such as the mouse. It is estimated that up to 50% of bovine conceptus loss occurs during the second and third week of pregnancy (1, 2), a period when a viable blastocyst undergoes extensive cellular proliferation and changes from a spherical shape to an elongated, filamentous form in preparation for implantation (3). During this period, three cellular lineages form in the hatched bovine blastocyst, epiblast, hypoblast, and trophectoderm. As seen in all mammalian species, these lineages will give rise to the embryonic disc developing into three germ layers, yolk sac after implantation, and the placenta upon differentiation, respectively. At the molecular level, this critical stage of development has only been characterized in the mouse model (4, 5), and more recently in non-human primates (6), and human embryo extended culture models (7, 8).

Peri-implantation development exhibits wide variation between species in relation to the duration of peri-attachment periods, the development and orientation of the extraembryonic tissues, and the implantation strategies (9). In the mouse, implantation occurs soon after blastocyst hatching from the zona pellucida. Within only a few embryonic days it extends from implantation to placentation, where several dramatic and concurrent events occur, making it difficult to study the molecular and cellular changes during this time period in rodents. Studying peri-implantation development in humans is also problematic as embryos embed into maternal tissues after the blastocyst stage, and ethical issues limit the scope of possible research. However, in ruminants, like cattle, implantation of blastocysts is preceded by a period of rapid growth and elongation. This peri-implantation period is prolonged compared to rodents, and is similar to that seen with human embryos. In this regard, the bovine is recognized as a highly informative model for human embryo development (10–13).

However, the cell types in the developing peri-implantation embryo and molecular mechanisms governing the embryo elongation in ruminants remain unexplored. To fill this knowledge gap, we collected bovine embryos at day 12, 14, 16, and 18, and established a comprehensive single-cell transcriptomic landscapes of peri-implantation development. Using bioinformatics analyses, we define the development of three major cell lineages (trophoblast, hypoblast, and embryonic disc) and their gene expression dynamics throughout peri-implantation development. Together with a comparative analysis of bovine peri-implantation trophoblasts and mature day 195 placental trophoblasts (14), we define and present a previously undefined trophoblast lineage. We also analyze cell-cell interaction signaling underling embryonic and extraembryonic cells interaction to ensure proper early development. This foundational information is useful to advance future efforts to understanding peri-implantation biology and causes of early pregnancy failure in the cattle.

## Results

### Construction of a single-cell transcriptomic census during bovine peri-implantation embryo development

To identify cell types and trajectories that lay the foundation for understanding bovine peri-implantation development, we performed single cell RNA sequencing (scRNA-seq) using the 10X Genomics Chromium platform (**Figure 1A**). We sequenced RNA of individual cells from bovine peri-implantation embryos at day 12 (pooled ten embryos to increase the cell populations), 14, 16, and 18 with biological replicates (**Supplementary figure 1A-D**). A total of 139,174 single cells from all peri-implantation stages were analyzed. Joint uniform manifold and projection (UMAPs) and clustering analysis revealed 10 distinct cell clusters for all cells from each of individual developmental stages (**Figure 1B-E, supplementary figure 1E-H**). To annotate the identities of cell clusters, we analyzed the database of known cell lineage markers in bovine (15–19), humans and mouse (20–22) and selected the markers that were detected in our single cell transcriptomes of bovine peri-implantation embryos (**Supplementary Dataset 1**). We identified *VIM*, *NANOG, MLY4, SLIT2, ACTA2, COL1A2,* and *BMP4* as marker genes of embryonic disc (ED), *SOX17, GATA4, FST, FN1, CDH2* and *CLU* as marker genes of hypoblast (HB), and *FURIN, IFNT, SFN, DAB2, PAG2* and *PTGS2* as trophoblast cell markers (**Figure 1G**). Using these markers, we captured three apparent major cell types in all four developmental stages, with 609 cells as ED, 21,283 cells as HB, and 117,282 cells as TB cell lineages (**Figure 1B-E**). Clustering analysis further revealed three subtypes of hypoblast cells and six subtypes of trophoblast cells (**Figure 1B-F**). As expected, the majority of cells analyzed were trophoblast cells due to the dramatic trophectoderm (TE) elongation during bovine peri-implantation development. We found a clearly developmental progression of cell lineage transition as evidenced by alternations of cell clusters between early (day 12 and 14) and later (day 16 and 18) peri-implantation stages, particularly trophoblast cell development (**Figure 1B-E**).

**Figure 1.**
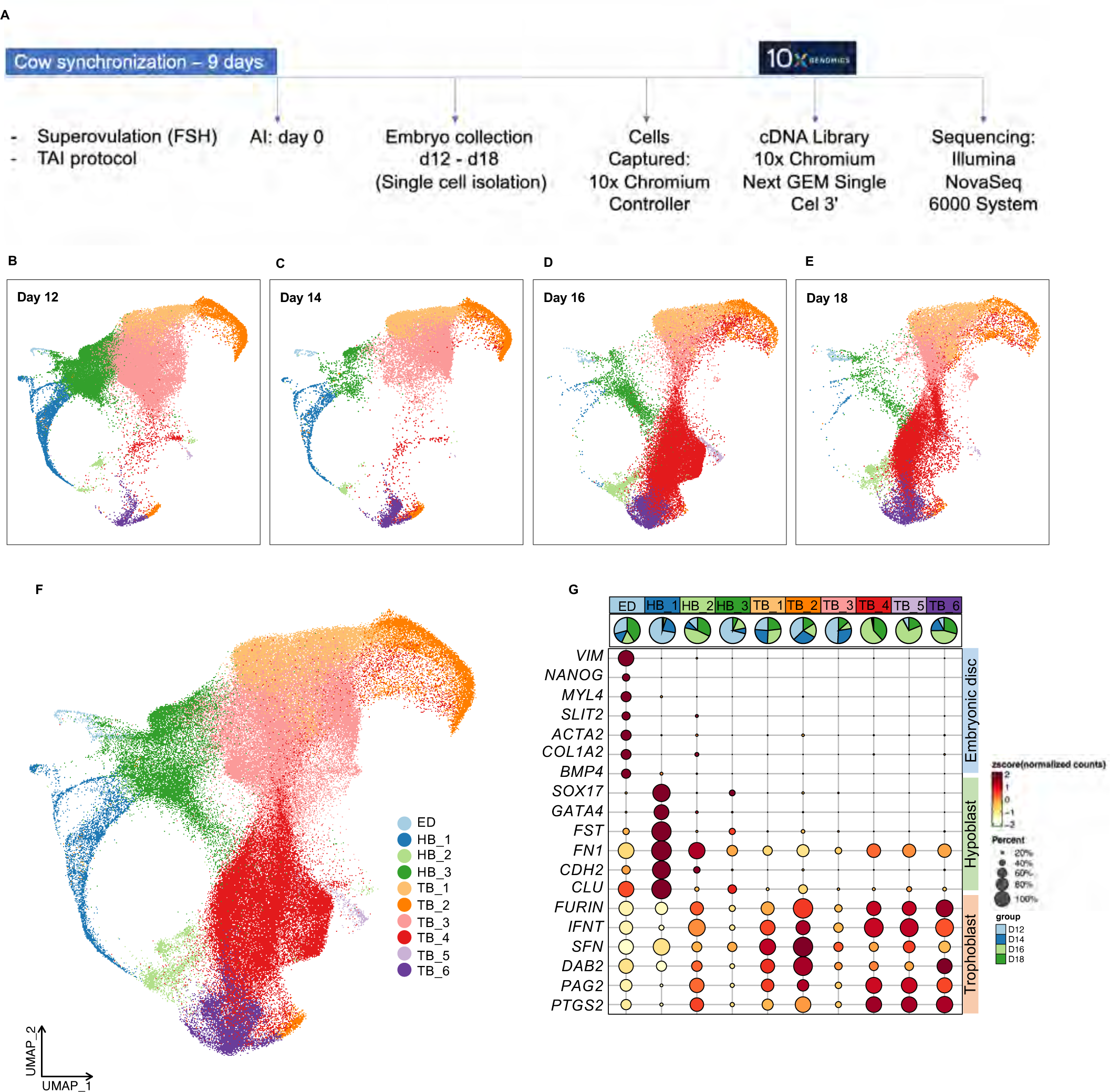
Single cell RNA-seq (scRNA-seq) analysis of bovine peri-implantation embryo development. **(A).** Diagram of cow synchronization protocol, embryo collection and single-cell isolation and scRNA-seq procedures with 10x genomics approaches. **(B-E).** Joint uniform manifold and projection (UMAP) analysis of transcriptomes of cell lineages from bovine peri-implantation embryos at day 12 **(B),** 14 **(C)**, 16 **(D)** and 18 **(E)**, dynamic lineage developmental progress is observed from day 12 through day 18. **(F).** UMAP clustering analysis of all cells captured from different developmental stages of bovine peri-implantation embryos. UMAP of integrated samples revealing 10 distinct cell types identified as embryonic disc (ED), hypoblast (HB) and different types of trophoblast cells (TB) cells. **(G).** Dot plot representing the expression of gene markers for ED, HB and TB development. Dot sizes represent the percentage of cells in the cluster expressing the gene marker, color gradient represents the level of expression from high (red) to low (yellow) and pie chart represent the stage present in the cluster.

### Development of embryonic disc during bovine peri-implantation development

ED development is one of the major events during bovine embryo elongation (23). After epiblast and hypoblast segregation in the inner cell mass (ICM) at day 9 post fertilization, the epiblast lineage further differentiates and forms the embryonic disc, which will contribute to the fetus after implantation (24). In the merged cell populations from day 12 to day 18, ED cells were clustered into two sub-clusters that were clearly separated in UMAP (**Figure 2A, top on the right panel**). Intriguingly, group of cells in the cluster marked as blue were exclusively from day 16 and day 18 embryos (ED late), while cells in the other cluster (red) were from day 12 and day 14 embryos (ED early) (**Figure 2A, bottom on the right panel**). By further investigating the expression patterns of three germ layer markers (endoderm: *AFP, SOX17, HHEX, FOXA2* (16, 25), mesoderm: *VIM, BMP4, ROR2, SOX6, FOXF1 and MSX1* (16, 26), and ectoderm: *NES, PARD6DB, MEIS* (27, 28), only cells in the ED late cluster showed the anticipated increase of those marker genes except for common endoderm markers, indicating mesoderm and ectoderm form by embryonic day 16 in bovine (**Figure 2A, left panel**). To explore the initiation of germ layer development, we performed clustering analysis only on the cells belonging to the embryonic germ layers (ED late, **Figure 2B, left panel)**. These cells were divided into two new sub-clusters (ED late_1 and ED late_2), which highly expressed mesoderm and ectoderm markers, respectively (**Figure 2B, right panel**). Together, these results indicate that germ layer development starts from day 16 after fertilization in bovine, where the formation and segregation of mesoderm and ectoderm lineages emerge, followed by the endoderm layer arising after day 18.

**Figure 2.**
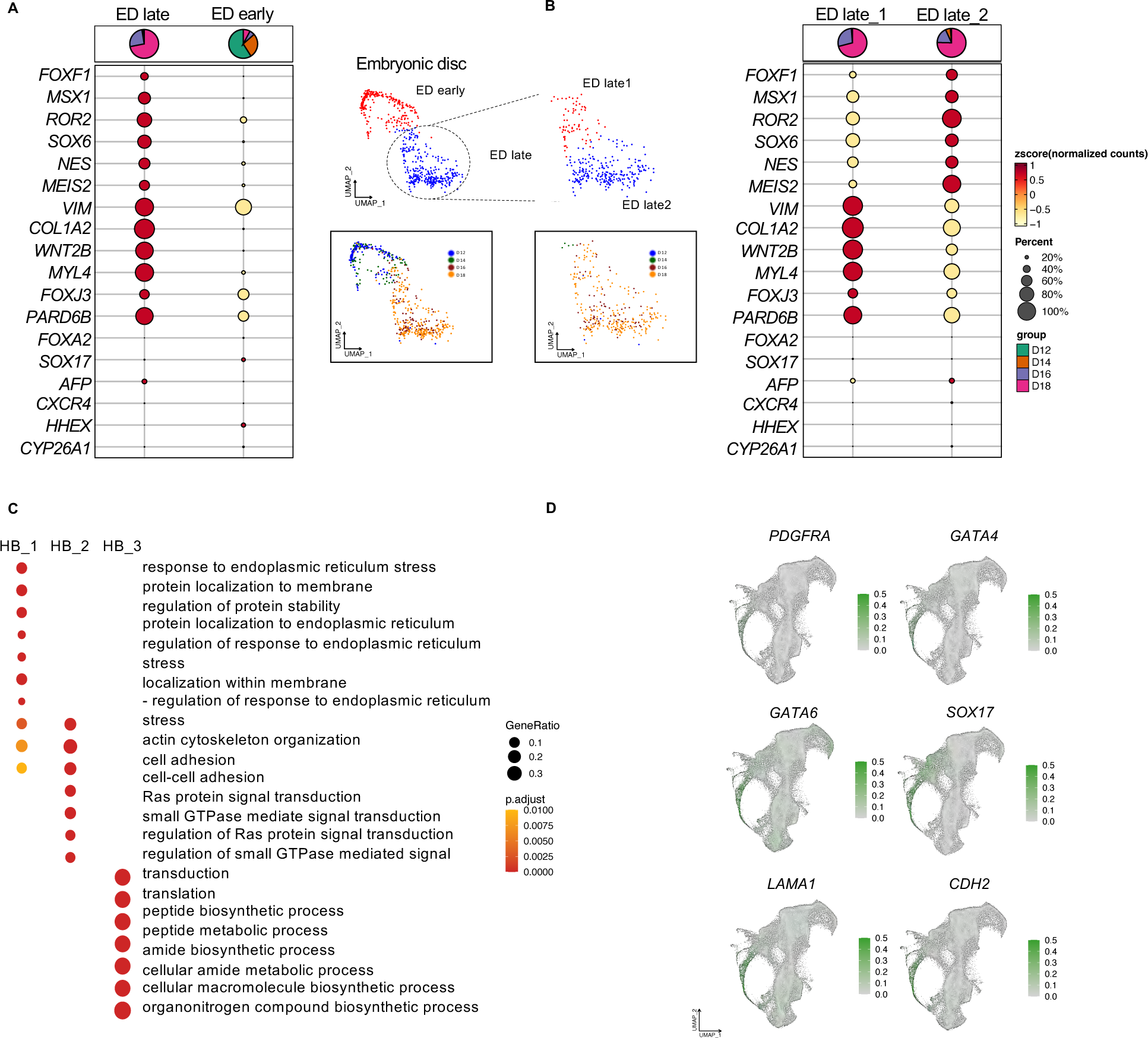
Identification of embryonic disc, germ layers and hypoblast lineages during embryo elongation. **(A).** Dotplot (left panel) and clustering (right panel) analysis of embryonic disc (ED) cell lineages. Two sub-clusters of ED were revealed (ED early and ED late). Highlighted area in blue were cells exclusively from day 16 and 18 (ED late) and red area with cells from day 12 and 14 (ED early). ED late highly expressed mesoderm and ectoderm markers (red dots). **(B).** Re-clustering analysis of ED late cell lineages. Two additional sub cell types were revealed (ED late_1 and 2). Dotplot analysis of expression of markers for ectoderm, mesoderm and endoderm in the ED late_1 and 2 cells. Dot sizes represent the percentage of cells in the cluster expressing the gene, color gradient represents the level of expression from high (red) to low (yellow) and pie chart represent the stage present in the cluster. **(C).** Dotplot showing the top represented GOs of genes specifically expressed in each of the hypoblast sub-lineages. **(D).** UMAPs showing the expression levels of common trophoblast markers (PDGFRA, GATA4, *GATA6, SOX17, LAMA1 and CDH2)* among hypoblast clusters. The color gradient from gray to green at the right refers to the gene expression level (high expression = green).

### Development of hypoblast during bovine peri-implantation development

Primitive endoderm (PE) or hypoblast (HB), which gives rise to yolk sac after implantation, is critical to support early conceptus development (29). We next characterized hypoblast development during bovine peri-implantation development. Three subtypes of hypoblasts were identified and showed distinct characteristics, 1) the majority of HB cells were of the HB_1 subtype in embryos at day 12 and 14 (**blue, Figure 1B-E**), HB_2 cells were present across all stages with increased cell populations during development from day 12 to 18 (**light green, Figure 1B-E),** on the contrary, HB_3 cells had decreased populations from day 12 to day 18 and clustered more closely to HB_2 cells as development progressed (**dark green, Figure 1B-E**); 2) functional gene ontology (GO) analysis of the highly expressed genes in each of the hypoblast cell subtypes revealed a significant enrichment in expression of genes related to response to endoplasmic reticulum stress, protein localization to membrane, regulation of protein stability in HB_1 cells, actin cytoskeleton organization, cell adhesion, ras protein signal transduction in HB_2 cells, and finally translation, biosynthetic process, and metabolic process in HB_3 cells (**Figure 2C**); 3) well known hypoblast lineage markers showed unique patterns between the subtypes, i.e., HB_1 cells were marked by *PDGFRA* and *GATA4,* HB_1 and 3 were positive for *GATA6* and *SOX17,* and LAMA1 and CDH2 were enriched in HB_1 and 2 cells (**Figure 2D**). This is consistent with the notion that these lineage markers contribute to conserved hypoblast lineage segregation in different mammalian species (17, 30). 4) we characterized the developmental progression of the three hypoblast subtypes by trajectory analysis and found they are originated as HB_1, progressed towards HB_3 and finally to HB_2 (**Supplementary Figure 3B**). The presence of HB_1 and 3 during early stages following by HB_2 present at later stages, suggesting a coordinated development of hypoblast lineages during bovine embryo development.

### Dynamics of trophoblast lineage development during bovine peri-implantation development

Trophectoderm elongation is an unique process in ruminants, where undifferentiated trophectoderm cells, or trophoblast progenitor cells will differentiate to mononucleated or uninucleate trophoblast cells (UNC) to drive embryo elongation and secrete IFNT, a signal for maternal fetal recognition (31, 32), and a subset will subsequently differentiate into binucleate cells (BNC) (33, 34), in preparation for attachment with maternal endometrium (3, 34). Most studies provide abundant data concerning the trophectoderm of blastocysts (17, 18) or trophoblasts after placentation (34), however, the trophoblast cell fate during the bovine peri-implantation period remain poorly understood. Using 14 widely accepted marker genes for bovine trophoblast cell lineages (16, 35), we were able to classify six trophoblast subtypes into two major lineages, the first with proliferative potential highly expressing *ASCL2*, *CDX2 and RAB25,* thus were defined as UNC (TB_1, 2, and 3), and a second subtype with increased expression of trophoblast markers including *IFNT*, *PTGS2* and *SSLP1* but not binucleate cell markers, and were therefore deemed as pre-BNC (TB_4, 5, and 6) (**Figure 1B-G**, **Figure 3A-B**). Our immunostaining analysis further confirmed the presence of trophoblast marker *PTGS2* and the absence of mature BNCs in peri-implantation embryos from day 12 to 18 (**Supplementary Figure 4C-D**), which is consistent with previous observation that BNC begins to appear by day 20 of pregnancy (33, 34). During trophoblast development, Day 12 and 14 TB were similar but they were very distinct from those at day 16 and 18, demonstrating a dramatic change of trophoblast dynamics from UNCs in day 12 and 14 to more emphasis on the pre-BNCs in day 16 and 18 (**Figure 1B-E**). This was further confirmed by the pseudotime trajectory analysis showing trophoblast development starts with TB_1, 2, 3 (UNC that are towards to the right edge of tree) and progresses towards TB_4, 5, 6 (pre-BNCs that enrich on the left edge of tree) (**Figure 3C**).

**Figure 3.**
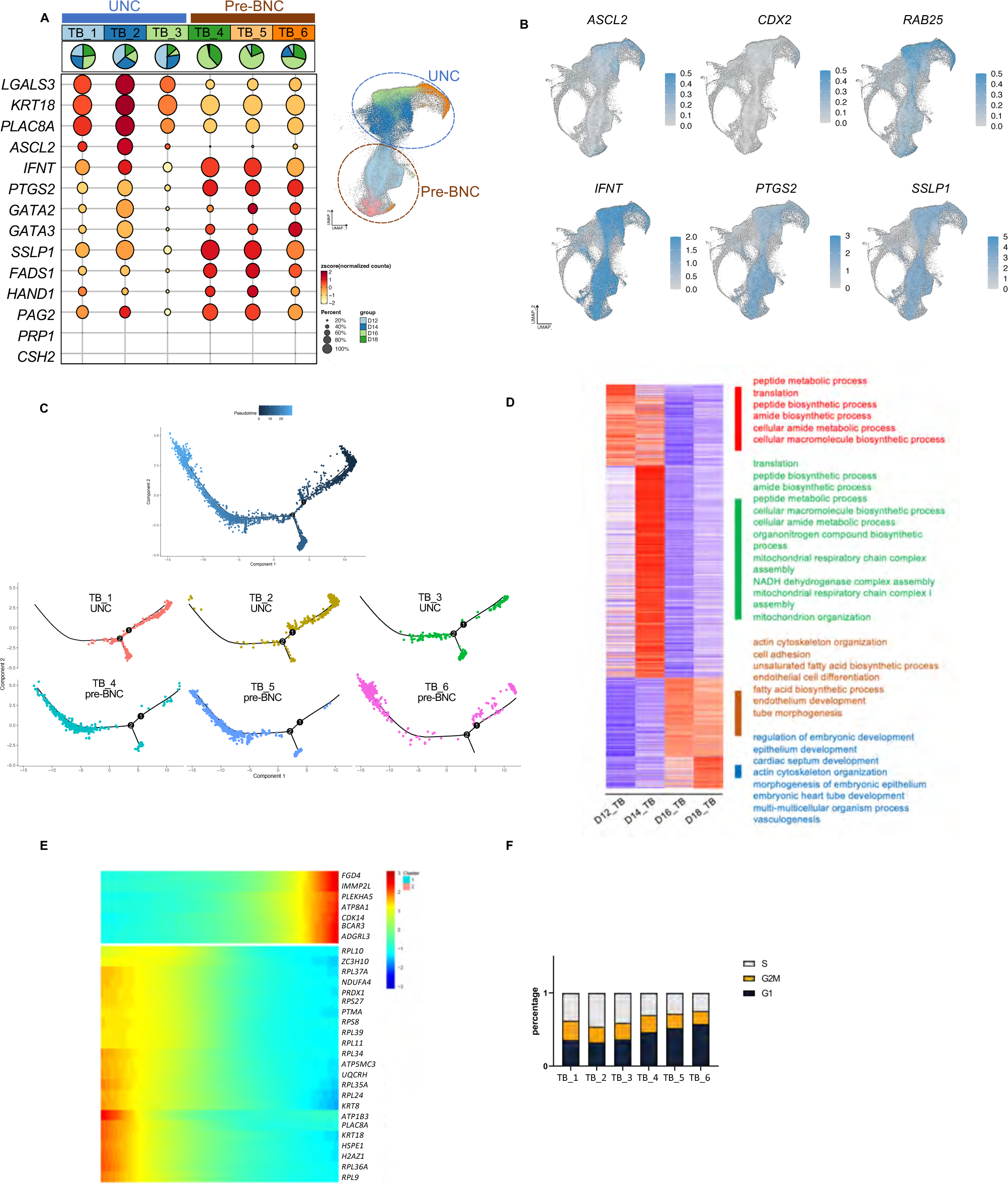
Dynamics of trophoblast lineage development during bovine peri-implantation development. **(A).** Dotplot (left panel) presenting common trophoblast markers in trophoblast clusters and classification of UNC and pre-BNC cells. Dot sizes represent the percentage of cells in the cluster expressing the gene, color gradient represents the level of expression from high (red) to low (yellow) and pie chart represent the stage present in the cluster. In the right panel, UMAP of trophoblast cell lineages divided in UNCs in blue circle (formed by TB_1, 2, and 3) and pre-BNC in brown (formed by TB_4, 5 and 6) **(B).** UMAPs showing the expression levels of *ASCL2, CDX2, RAB25, IFNT, PTGS2 and SSLP1* (common trophoblast markers). Diverse expression level among trophoblast clusters (*ASCL2*, *CDX2 and RAB25)* expressed in UNC; (IFNT, PTGS2 and SSLP1) more expressed in pre-BNC cells. **(C).** Pseudotime colored in gradient from dark to light blue. Two start points of development was identified in the right edge by dark blue (top panel). In the bottom panel, the distribution of clusters demonstrated the development of TB_1, 2 and 3 (UNC) into TB_4, 5, and 6 (pre-BNC). **(D).** Heat map showing top enriched pathways from genes specifically expressed in each developmental stage of trophoblasts. **(E).** Heatmap showing scaled expression of dynamic genes along Pseudotime of trophoblast development. The color bar represents the z-score distribution from -3 (blue) to 3 (red). **(F).** Cell cycle composition of TB clusters confirming higher proliferative status of pre-mature BNC.

To understand the biological function of trophoblast sub-lineages during bovine elongation, we first analyzed the trophoblast stage specific genes that corresponded to different peri-implantation stages (**Figure 3D**). Interestingly, analysis of the functions of these stage specific genes revealed a sequential progression of trophoblast stage-specific core gene networks. It migrated from peptide metabolic processes, translation, and biosynthetic processes in day 12, to regulation of translation, metabolic processes and mitochondrial function in day 14, to actin cytoskeleton organization, cell adhesion, and fatty acid metabolism processes in day 16, and finally to regulation of embryonic development, epithelium, tube, tissue, blood vessel and vasculature development in day 18 (**Figure 3D**). Such coordinated changes of functional pathways further confirmed trophoblast cell development transitions from UNC to pre-BNC cells and are reflective of the general lack of knowledge concerning trophoblast lineage identities and gene expression patterns during this critical period of development in cattle. Second, we explored the specific genes with enriched expression in the transition of the two major trophoblast lineages. It was found that several trophoblast marker genes and genes related to ribosome activity were highly expressed in UNC including *KRT8, KRT18, PLAC8A, H2AZ*, and RPL (Ribosomal Protein Large) subunit gene family (**Figure 3E**). However, genes highly expressed in pre-BNC were closely related to tumorigenesis, such as *BCAR3* (36), *FGD4* (*37*), and *PLEKHA5* (*38*), suggesting pre-BNC might be necessary for embryo elongation, attachment and implantation (9, 34). Third, we analyzed cell cycle composition of separate trophoblast clusters and found a higher proliferative status in pre-BNC cells. (**Figure 3F)**. Finally, a list of highly expressed genes were identified in a trophoblast cell subtype-specific manner (**Supplementary Figure 4A**). The genes with most dynamic changes during trophoblast development included *ADAMTS1, AHSG, ATP5PO, CSTB, FETUB, LPP, PTTG1IP* and *TP63* (**Supplementary figure 4B**).

### Comparative analysis of bovine peri-implantation trophoblasts and mature day 195 placental trophoblasts

With the publicly available single cell transcriptomes of mature trophoblasts from bovine day 195 placenta (14), we sought to construct a transcriptomic road map of bovine trophoblast differentiation. The peri-implantation trophoblasts and mature placenta trophoblasts were grouped together (distinct from ED and HP, data not shown) and formed 10 different clusters **(Figure 4A left**), clearly separated by their developmental stages (**Figure 4A right**). The marker gene analysis further confirmed the absence of binuclear cells in the bovine peri-implantation embryos till day 18 (**Figure 4B**), suggesting that the newly identified primitive trophoblast cells (pre-BNCs) are responsible for pregnancy maintenance in bovine prior to the time when binucleate cells emerges. Trajectory analysis was performed showing peri-implantation trophoblast cells develop into mature placenta trophoblasts as expected but were separated by the uncharacterized trophoblast cell lineages (**Figure 4C**). In addition, we identified the genes with most dynamic changes in the transition between trophoblasts from peri-implantation embryos and mature placentas included 1) *PDXK, FETUB, LPP* and *AHSG* have high expression specifically in peri-implantation trophoblasts although they are very dynamic between UNC and pre-BNC cells (**Figure 4D**), indicating their essential functions in the respective trophoblast lineages; and 2) *EIF4A2* and *TMEM50B* that have increased levels in mature placenta trophoblast compared to UNCs and pre-BNCs (**Figure 4D right**). Interestingly, all of these genes have their known functions in cell proliferation and tumorigenesis (39–42), suggesting that, similar to human trophoblast development (8), the proliferation and migration/invasion activity are two important functional indicators to classify bovine trophoblast cell lineages.

**Figure 4.**
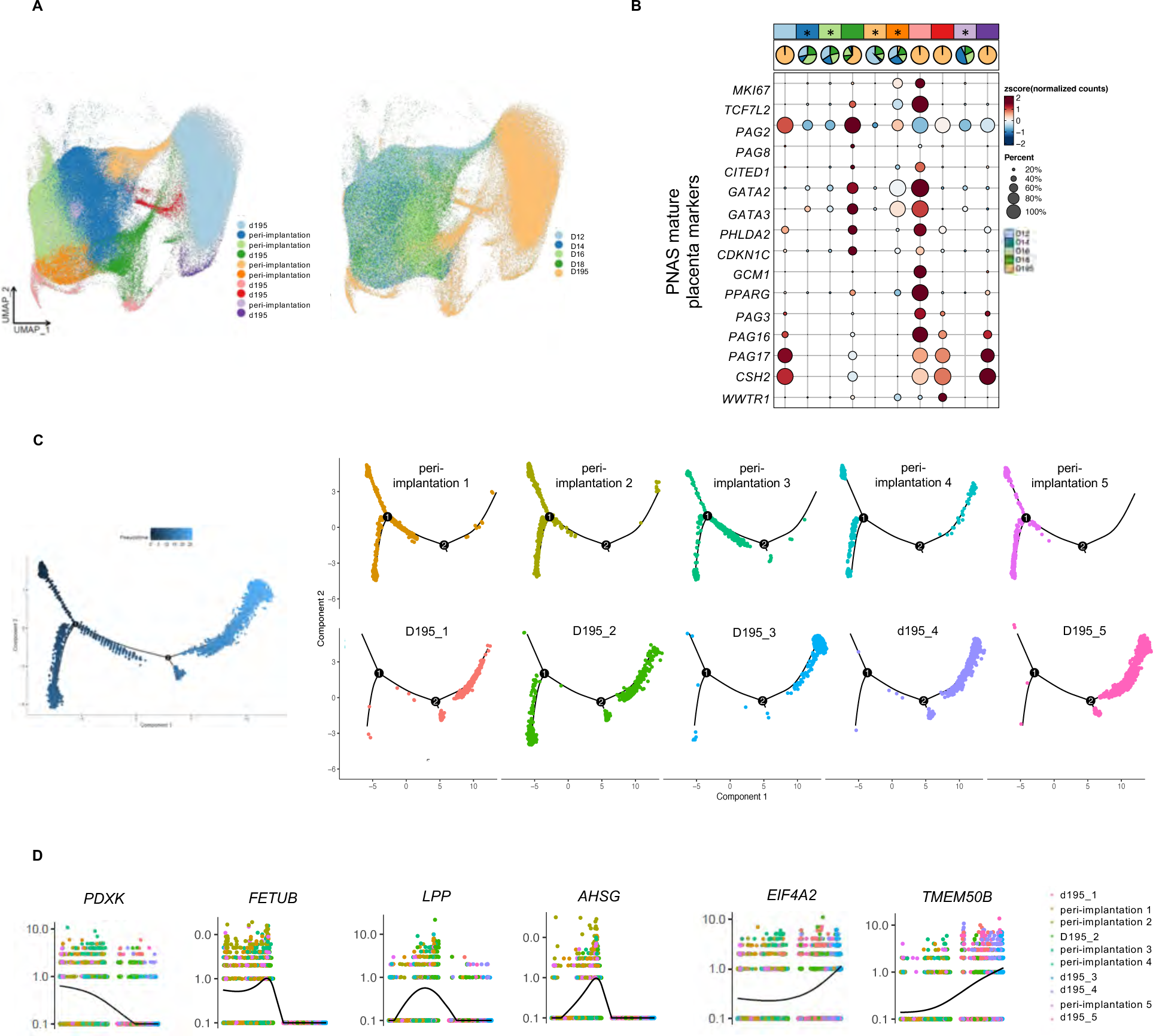
Comparative analysis of bovine peri-implantation trophoblasts and mature day 195 placental trophoblasts. **(A).** UMAPs of integrated single cell trophoblast lineages from peri-implantation and mature placenta. All identified trophoblast clusters from both datasets were presented on the left panel, trophoblast clusters were presented by developmental stages on the right panel. **(B).** Dotplots representing the expression levels of common trophoblast markers from mature placenta trophoblasts. Clusters that are exclusively from peri-implantation stages were marked by “*” and had lower or no expression of BNC markers. Dot sizes represents the percentage of cells in the cluster expressing the gene, color gradient from red to yellow represents the level of expression from high to low and pie chart represent the stage present in the cluster. **(C).** Pseudotime trajectory analysis of trophoblast development of TB peri-implantation stages and TB mature placenta colored in gradient from dark to light blue. Two start points of development was identified in the left edge by dark blue (left panel). On the right panel, the distribution of clusters confirmed the development of peri-implantation trophoblast into trophoblast from mature placenta trophoblast. **(D).** Top six differentially expressed genes (*FETUB, PDXK, LPP, AHSG, EIF4A2* and *TMEM50B*) that had most dynamic expression changes between peri-implantation trophoblasts and mature placenta trophoblasts. The black line is the mean fitted expression level across the sample.

### Identification of the transcriptional factors and novel lineage markers during bovine peri-implantation development

Given that most known transcriptional factors (TFs) that are essential developmental regulators, are limited to pre-implantation embryos (43, 44), here we identified transcriptional factors (TFs) during bovine peri-implantation embryos and explored the key regulators directing the development of specific cell lineages (**Supplementary Figure 2A**). For example, important mediators for trophoblast stem cell self-renewal (*CDX2, ESRRB, GATA2, GATA3, TFAP2A*, and *TFAP2C*) (45), epiblast development (*TEAD2*) (46), and primitive endoderm (*GATA4, GATA6, SOX17,* and *TBX3*) (47) were prioritized in the bovine peri-implantation development (**Supplementary Figure 2A**). We also identified little known regulators for lineage specification including *PRDM6, TGIF1* and *HNF1B* (**Supplementary Figure 2A**).

Since most of the early cell lineages markers have been studied in mouse early development, we next sought to identify the novel markers for the bovine early cell lineages. First, we identified novel markers associated with identified ED (**Supplementary Figure 5A**). These highly expressed genes specific in the bovine ED lineages included *CDC42EP5*, a regulator of cytoskeleton organization and migration (48), *PRTG*, that is essential for mesoderm and nervous tissue development (49) and *PLTP*, a mediator of lipoprotein metabolism and transport (50) (**Supplementary Figure 2B**). Next, we analyzed the genes that are significantly upregulated in a cell type-specific manner in hypoblast lineages and prioritized the top 10 novel markers of three hypoblast cell subtypes (**Supplementary Figure 3A, 5A**) including *CTSV* and *RASGRF2* (**Supplementary figure 2C**). Finally, we examined novel markers for trophoblast cell lineages and found that *CFAP54* and *TMEM86A* had a significant high expression in UNC cells, while *PLEKHA5, SATB2* and *RAPGEF2* marked pre-BNC cells. (**Supplementary Figure 2D, 5A**). A deeper investigation of these novel markers could lead to elucidate new insights into mechanisms that facilitate bovine implantation.

### Embryonic and extraembryonic cell-cell interactions during bovine peri-implantation development

Faithful embryogenesis and success of pregnancy establishment require a precise coordination between embryonic and extraembryonic lineages (51). Here we sought to identify the cell-cell interaction signaling between lineages contributing to embryonic and extraembryonic tissues in bovine. We explored the signaling interactions (**Figure 5A**) and identified ligand and receptor pairs (**Figure 5B**) among lineages using Cell Chat analysis. Based on the number of signaling interactions from each lineages, we found that TB_1, 3 and hypo_3 lineages work independently from other cells with less outgoing and incoming signaling (**Figure 5B**). Conversely, Hypo_1 was shown to be an interactive lineage, with the most sender and receiver signaling. Additionally, HB_1 received massive signals from TB_2 (UNCs) and sent most of the signals to TB_6 (pre-BNCs) (**Figure 5B**), suggesting that hypo_1 could be an important mediator for trophoblast differentiation and thus promotes embryo elongation.

**Figure 5.**
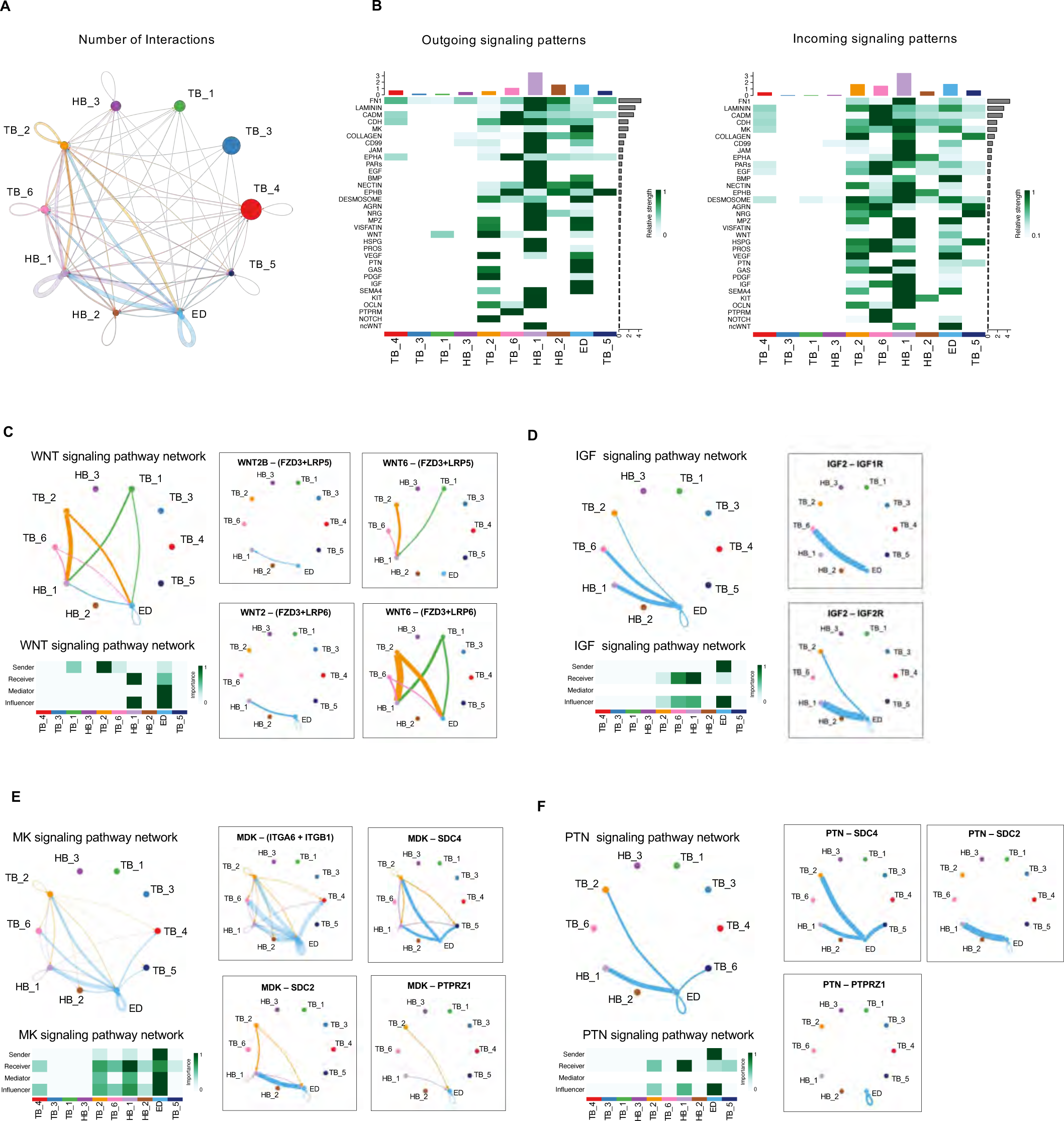
Embryonic and extraembryonic cell-cell interactions during bovine peri-implantation development. **(A).** Circle plot analysis showing the significant cell–cell interaction among different cell lineages. Arrows and edge color indicate direction (ligand: receptor), the circle size represents the number of cells and edge thickness indicates the communication probability. **(B).** Heatmap showing the identified pairs of ligand and receptor signaling in the embryonic and extra-embryonic cell lineages of bovine peri-implantation embryo. Outgoing signaling were presented on the left panel, incoming signaling were presented on the right panel. The color gradient on the left represents the relative signaling strength of the signaling pathway across clusters. **(C, D and E).** On the left panel, circle plot showing the intercellular communication network for WNT, IGF, MK and PTN signaling and heatmap showing the relative importance and contribution of each cell lineage to the overall communication network. On the right panel, Circle plot showing the identified ligands and respective receptors in each signaling.

Of note, two well-known signaling pathways, WNT and IGF, were found to be outgoing signaling in UNC (**Figure 5C**), and an outgoing signaling in ED (**Figure 5D)**, respectively. It is noteworthy that IGF is essential for fetus development and growth (52), while WNT signaling is a crucial factor affecting both embryonic and extraembryonic stem cell maintenance (53, 54). Additionally, we found ED was an important sender of MK and PTN signaling that have important function in cell proliferation, migration and self-renew (55). Interestingly, both of them shared the same receptors (PTPRZ1 and SDC2/4) in both hypoblast and trophoblast lineages (**Figure 5E, F**), suggesting hypoblast and ED mediate the development of extraembryonic lineage development.

Together, the identification of these cell-cell interaction molecules provides candidate regulators for further mechanistic studies underly how embryonic and extraembryonic cells interact to ensure proper early development in bovine.

## Discussion

Peri-implantation development is a critical period when most pregnancies fail, yet is the least studied process during mammalian development. In the bovine, peri-implantation is defined by embryo elongation. During this process, the blastocyst cells proliferate massively, the initially un-limited potential of the epiblast is restricted and shaped by a combination of changes in cell lineage composition. Additionally, gene expression, cell-cell interactions, and physical forces - toward defined germ layers and cell types, as well as the trophoblast progenitors differentiating to form primitive placenta and initiate maternal fetal recognition (31, 32) all work to establish the successfully pregnancy. Here, we have provided a single cell transcriptomic wide characterization of these cellular and molecular events accompanying the bovine embryo elongation. The datasets, particularly when mined further and integrated with epigenome information, are expected to greatly expand our understanding of the gene regulation mechanisms governing bovine peri-implantation embryo development, which will provide some insight into what can potentially go wrong in the pregnancies that fail during this period.

Our study revealed the timing and cell types emerging in a coordinate fashion during bovine early development and observed some surprises. First, epiblast, later embryonic disc developed into mesoderm and ectoderm in embryo day 16 and 18 while the endoderm emerged much later. This is quite interesting as it is in contrast with the mouse, where the declining of epiblast cells is followed by mesoderm and endoderm lineage development and ectoderm development one day later (56). Second, our analysis identified a previously unrecognized primitive trophoblast cell lineage, termed as pre-BNCs and confirmed the absence of the binuclear cells in a peri-implantation stage embryo. Third, trophoblast cell development is very dynamic, trophoblast cells at day 12 and 14 represent UNCs, while a major shift occurs at day 16, when pre-BNC cells become dominant. This dramatic change is also coordinate with the embryo’s dramatic elongation in size from an elongated form at day 14 to a filamentous form at day 16. Interestingly, the embryo changes from a spherical shape at day 12 to elongated form at day 14, however, day 12 and day 14 embryos have very similar cell composition and transcriptomes, suggesting the bovine embryo elongation may not be driven by embryo internal genetic factors.

As expected, most of the cells analyzed in the bovine peri-implantation embryo are trophoblast cell lineages. Our analysis identified 6 different sub-lineages of trophoblast cells, classified into two categories as UNC and pre-BNC cells. Three sub-types of pre-BNC cells exhibited very interesting transcriptomic features. First, genes highly expressed in TB_4 were related with epithelium, endothelial development and establishment of endothelial barrier. Second, Genes in TB_5 showed advanced functions with tube, blood vessel and vasculature development supporting the development and formation of the placenta vasculature. TB_5 gene clusters also exclusively expressed placenta developmental markers including *RXRA* and *CITED2*, which have been identified in mouse and human as syncytiotrophoblast and invasive extra villous respectively (22, 57). Third, TB_6 was associated with cytoskeleton organization, cell periphery and ras protein signal transduction. TB_6 cells also expressed exclusive important markers such as *AMOT, PPARG, and* WWTR1. AMOT plays an important role in TB segregation and function (58)*, PPARG* is essential for trophoblast differentiation and binuclear cell development (59), and WWTR1 is related with TB self-renew and EVT differentiation in human (60). In addition, many signaling pathways essential for mature trophoblast cells were enriched in our identified pre-BNC cells such as MAKP and VEGF important for placenta development and vascularization (61, 62), Hippo signaling is related with cell differentiation, fetal growth, and establishment of a connection between fetal and maternal circulation (63), mTOR in regulating placental growth (64), and Rap1 in cell polarity, cell interactions, cell adhesion and proliferation in early embryonic development (65). Many identified gene functions of pre-BNC cells have also been reported to be required for trophoblast differentiation such as cytoskeleton organization, cell differentiation, adhesion, periphery, migration and tight junction formation (34). Collectively, while binuclear cells are not present in the peri-implantation embryos, this newly identified pre-BNC cells have up-regulated machinery for the binuclear cells, and represent an important stage of trophoblast cell fate that are responsible for pregnancy maintenance in bovine prior to the time when the binucleate cells emerge. The identification of these progressive functions is also the first step to reveal the importance of pre-BNC cells in the formation of the functional placenta and maintenance of the pregnancy.

Perhaps most importantly, we identified novel markers of cell lineages emerging in the bovine peri-implantation development and the cell-cell interactions that mediate this unique embryo elongation process. While they are largely unexplored, a deeper understanding of their functional operation during these stages might facilitate our understanding of little-known bovine peri-implantation development.

In summary, our work has filled a significant knowledge gap in the study of lineage development over a period of rapid change of embryo elongation and provide foundational information to understanding peri-implantation biology and causes of early pregnancy failure in the cattle.

## Materials and Methods

### Animal care and use

Bovine peri-implantation embryos were collected from non-lactating, 3-year-old crossbreed (Bos taurus x Bos indicus) cows housed at the Reproductive Biological Center (RBC) at the School of Animal Sciences, Louisiana State University Agriculture Center (LSU AgCenter). The experiments were conducted under an animal use protocol (A2021-21) approved by the Louisiana State University Agricultural Center Institutional Animal Care and Use Committee.

### Cow synchronization

Cows were synchronized starting on day 0 with dominant follicle removal (DFR) followed by insertion of standard 7-day vaginal controlled internal drug release of progesterone (CIDR, Zoetis). On day 2 ovulation-inducing gonadotropin-release hormone (GnRH, Fertagyl, Merk Animal Health) was administered intramuscular (IM) injection. From day 4-7, follicle stimulating hormone (FSH, Folltropin, Vetoquinol) was administered twice a day in a decreasing dose. Upon CIDR removal on day 7, one dose of prostaglandin (Lutalyse, Zoetis) was administered in the morning and afternoon. 48 hours after CIDR removal another dose of GnRH was administered via IM injection and artificial insemination was procedure twice in a 12-hour interval.

### Embryos collection

Bovine peri-implantation embryos were collected at day 12, 14, 16 18 days after artificial insemination. Embryos were recovered by standard non-surgical flush with lactated ringer solution supplemented with 1% fetal bovine serum and washed with PBS before processing for single cell isolation. All cows were treated with prostaglandin (Lutalyse, Zoetis) after flushing.

### Single cell isolation

After embryo collection, fresh embryos were washed with PBS and placed in a 3% FBS in ice cold PBS. Embryos were centrifuged for 5 minutes at 400xg at 4°C. After supernatant aspiration, embryos were resuspended in 200μL of TrypLE and minced by scissors. 500μL of TrypLE were added and then embryos were incubated at 37°C in a shaker at 150rpm for 4-7 minutes depending on size of embryos. Samples were pipetted every 2 minutes to avoid large clumps. Dissociation was stopped with same volume of 3% FBS (700μL), and the suspensions were pass through 70μm cell strainer and centrifuged for 5 min at 400xg at 4°C. The cell pellet was resuspended with 0.04% BSA (volume depended on the size of cell pellet). Cell suspensions were filtered through 40μm cell strainer into a new 1.5mL Eppendorf tube. Cell viability and concentration were measured using a Countess Automated Cell Counter. The cells with viability at least 80% were proceeded with the 10x Genomics® Single Cell Protocol with a target of 10,000 cells per sample. Single cell libraries were prepared using 10x Chromium Next GEM Single Cel 3’ Reagent Kit v3.1 Dual Index followed manufacturer’s instructions. Libraries were sequenced with an Illumina Novaseq 6000 System (Novogene).

### Single-cell data pre-processing and clustering

To analyze 10X Genomics single-cell data, the base call files (BCL) were transferred to FASTQ files by using CellRanger (v.7.1.0) mkfastq with default parameters, followed by aligning to the most recent bovine reference genome downloaded from Ensembl database (UCD1.2.109), then the doublets were detected and removed from single cells by using Scrublet (0.2.3) with default parameters. The generated count matrices from all the samples were integrated by R package Seurat (4.3.0) utilizing canonical correlation analysis (CCA) with default parameters (https://satijalab.org/seurat/articles/get_started.html) (66). The data was scaled for linear dimension reduction and non-linear reduction using principal component analysis (PCA) and UMAP, respectively. The following clustering and visualization were performed by using the Seurat standard workflow with the parameters “dim=1:30” in “FindNeighbors” function and “resolution=0.2” in “FindClusters” function. The function “FindAllMarkers” in Seurat was used to identify differentially expressed genes in each defined cluster. The cutoff value to define the differentially expressed genes was p.adjust value < 0.05, and fold change > 0.25. The UMAP plots and bubble plots with marker genes were generated using “CellDimPlot” and “GroupHeatmap” functions in R package SCP (0.4.0) (https://github.com/zhanghao-njmu/SCP), respectively. Gene ontology (GO) and pathway analysis was performed using R package clusterProfiler (4.6.1), and the GO terms were presented by “dotplot” function in Seurat.

The raw FASTQ files and normalized read accounts per gene are available at Gene Expression Omnibus (GEO) (https://www.ncbi.nlm.nih.gov/geo/) under the accession number GSE234335.

### Constructing trajectory

Cell differentiation was inferred for trophoblast subtypes from peri-implantation embryos and from cotyledon part in Day195 of gestation using the Monocle 2 method (2.26.0) with default parameters (67). Because of the large amount of cell numbers, 1000 cells were randomly selected from each cluster and used for the following analysis. Integrated gene expression matrices with the smaller sample size from each subtype were exported into Monocle by constructing a CellDataSet. Genes detected less than 20 cells were removed and then the variable genes were defined by “differentialGeneTest” function, the top 1000 genes were used for cell ordering with the “setOrderingFilter” function. Dimensionality reduction was performed using the “DDRTree” reduction method in the reduceDimension step. The root of the pseudotime trajectory was assigned based on the time point of the development (clusters enriched at D12 was considered as root). Pseudotime related genes were defined by using “differentialGeneTest” function. Monocle 3 (1.3.1) was also used to construct the pseudotime trajectory from elongation embryos to D195 cotyledon cells with the default workflow steps (https://cole-trapnell-lab.github.io/monocle3/) (68).

### Single-cell regulatory network inference and clustering (SCENIC) analysis

We explored the transcription factor network inference by using the SCENIC R package (version 1.3.1, with the dependent packages RcisTarget 1.17.0, AUCell 1.20.1, and GENIE3 1.20.0) (69). Activity of the regulatory networks was evaluated by the standard workflow by using “runSCENIC_1_coexNetwork2modules”, “runSCENIC_2_createRegulons”, “runSCENIC_3_scoreCells”, and “runSCENIC_4_aucell_binarize” in a row. Then potential direct-binding targets (regulons) were explored based on motif analysis. Followed by that, based on the AUCell algorithm, SCENIC calculates each regulator’s activity and builds gene-expression rankings for each cell. To find the main transcription factors regulating bovine peri-implantation embryo development, the regulon activity was averaged. A regulon-group heat map was generated with pheatmap package in R.

### Cell–cell communication analysis

Potential cell–cell interactions based on the expression of known ligand–receptor pairs between different clusters were identified using CellChat (1.6.1) (70). Integrated gene expression matrices from all subtypes were exported from Seurat into CellChat by using “createCellChat” function, followed by preprocessing workflow steps including “dentifyOverExpressedGenes”, “identifyOverExpressedInteractions”, and “projectData” functions with default parameters. The cell-cell communication was then calculated by functions “computeCommunProb”, “filterCommunication”, “computeCommunProbPathway”, and “aggregateNet” in a row with default parameters. The significant intercellular signaling interactions for particular pathway families of molecules were performed with “netVisual” function. To determine the senders and receivers for specific pathways, the function netAnalysis_computeCentrality was applied on the netP data slot. The contribution of each cell subtype to enriched interaction pathways including both outgoing pattern and incoming pattern were visualized by using netAnalysis_signalingRole_heatmap function.

### Author contributions

Z.J designed and supervised research. G.S. performed most of the experiments. H.M. analyzed all genomic data. Y.W. performed single cell suspension. L.Z., E.C. and K.B. helped with embryo collection. G.S., H.M., and Z.J. interpreted data and assembled the results. G.S., H.M., and Z.J. wrote the manuscripts with inputs from all authors.

## Acknowledgements

We thank Dr. Joel Carter for his assistance with embryo flushing. This work was supported by the NIH Eunice Kennedy Shriver National Institute of Child Health and Human Development (R01HD102533) and USDA National Institute of Food and Agriculture (2019-67016-29863, W4171).

## Author contributions

Z.J designed and supervised research. G.S. performed most of the experiments. H.M. analyzed all genomic data. Y.W. performed single cell suspension. L.Z., E.C. and K.B.

## Supplementary Figures and Datasets

**Supplementary Figure 1.**
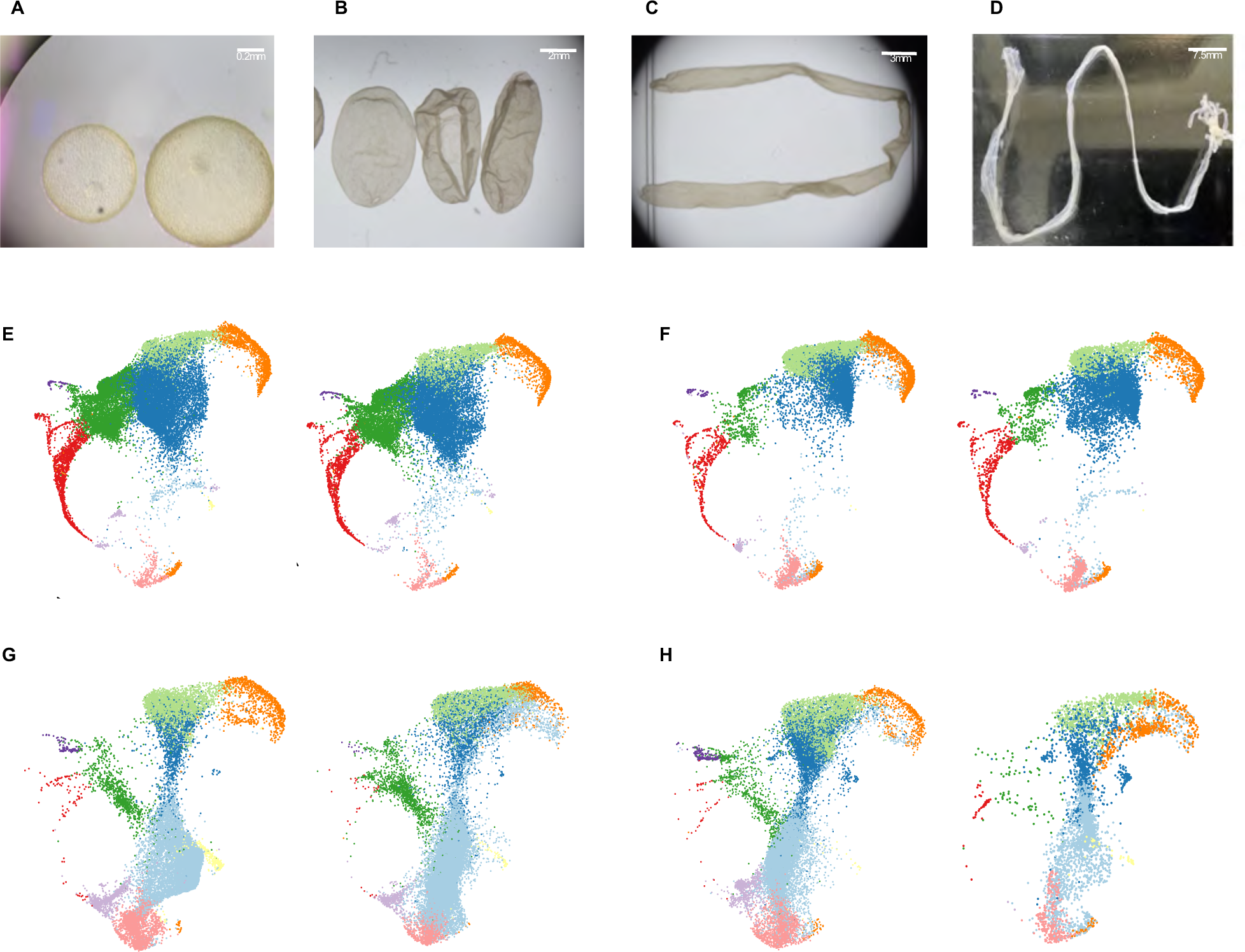
Represented bright field images of bovine peri-implantation embryos at day 12 (**A,** spherical shape), day 14, **(B,** ovoid shape), day 16 **(C,** filamentous shape), and day 18 **(D,** filamentous shape). **(E, F, G and H).** UMAP analysis of transcriptomes of cell lineages from bovine peri-implantation embryos at day 12 **(B),** 14 **(C)**, 16 **(D)** and 18 **(E)** with duplicates. Each color represents different lineage identified among the stages.

**Supplementary Figure 2.**
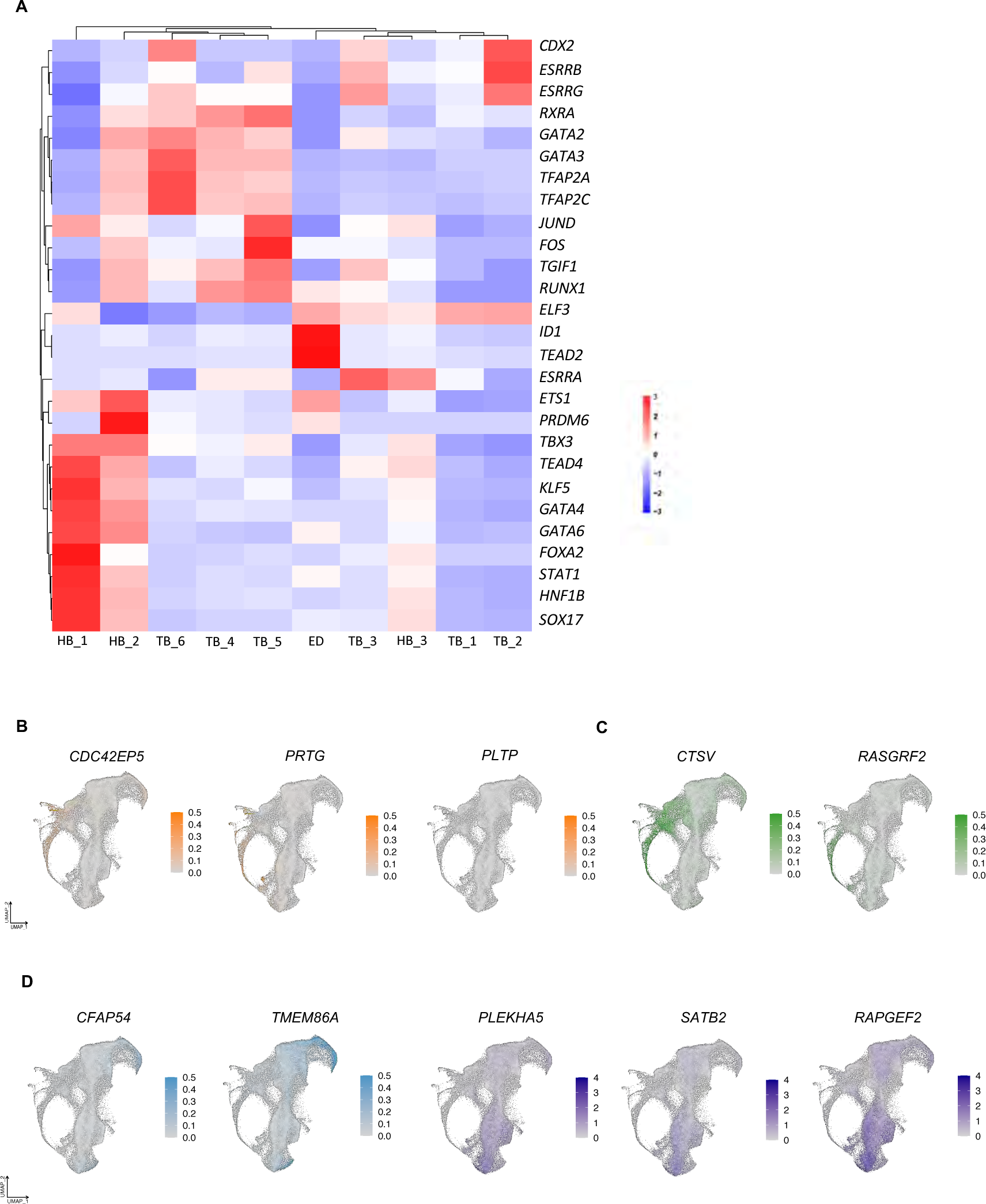
**(A).** Heatmap of top 10 transcription factors identified in bovine peri-implantation embryos. Each column represents a different lineage. The expression level is represented by the gradient color from blue to red. UMAPs showing the expression levels of identified novel makers in ED **(B, orange),** HB **(C, green)**, and UNC (**D, blue**) and pre-BNC cells (**D, purple**).

**Supplementary Figure 3.**
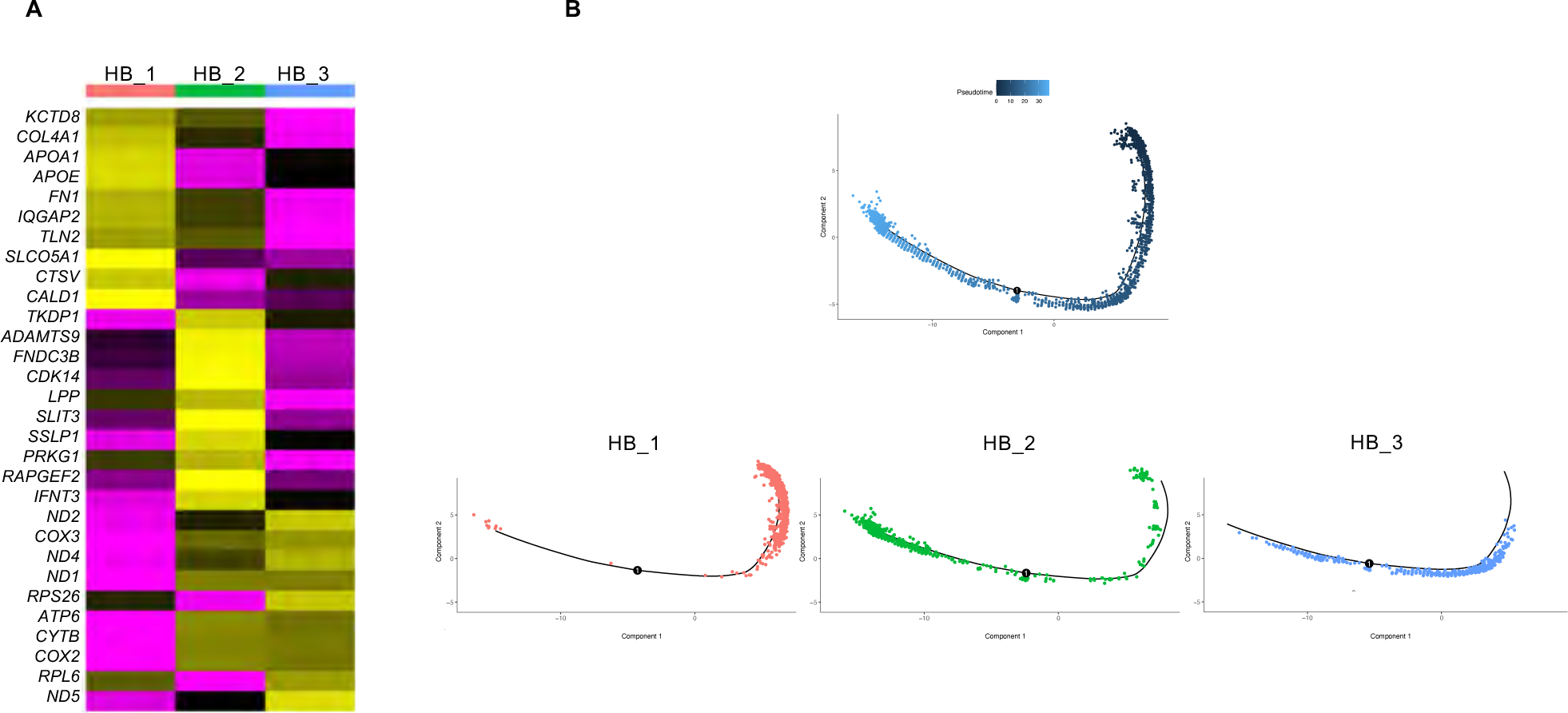
**(A).** Heatmap showing the top 10 gene markers identified in hypoblast clusters. **(B).** Pseudotime trajectory analysis of hypoblast clusters, colored in gradient from dark to light blue. The right edge (dark blue) represents the start point of development, followed by the distribution of hypoblast cells in each cluster confirming the development from hypo_1.

**Supplementary Figure 4.**
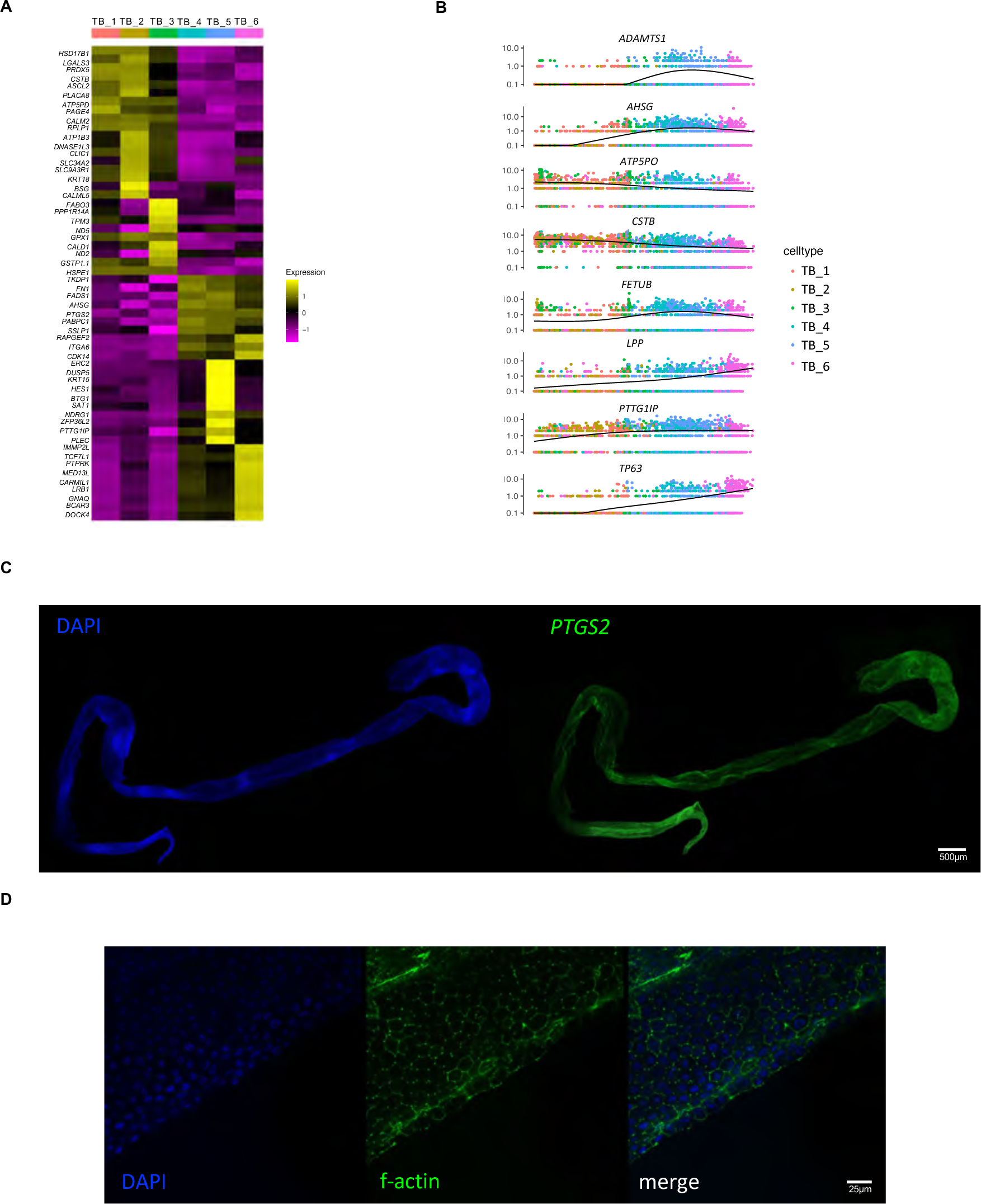
**(A).** Heatmap showing the top 10 genes markers identified in each trophoblast sub-lineages. **(B).** Dot plot showing the dynamic expression of trophoblast markers during TB development. (**C**). Immunostaining analysis of trophoblast cells in embryo at day 16. DAPI (blue), *PTGS2* (common trophoblast marker – green). (**D**). Immunostaining analysis of trophoblast cells in embryo at day 18. DAPI (blue), f-actin (green) and merged confirming absence of binuclear TB cells.

**Supplementary Figure 5. (A).** List of identified novel gene markers for ED, HB, and TB.

**Supplementary Dataset 1.** Differentially expressed genes among all 10 identified clusters.

| UNC | Pre-BNC | Hypoblast | Embryonic disc |
| --- | --- | --- | --- |
| PINLYP | ERC2 | CTSV | PLTP |
| CFAP54 | RBM47 | ADAMTS9 | PTMA |
| FXYD4 | PLEKHA5 | RASGRF2 | MS4A8 |
| TMEM86A | ADGRL3 | RCN1 | PSIP1 |
| SEC61G | RAPGEF2 | SKAP1 | CDC42EP5 |
| TOMM5 | RABPC1 | SPINT2 | PRTG |
| NDUFB1 | PTTG1LP | CASK | RORA |
| AMDHD2 | SATB2 | FOXP1 |  |
| C1QBP | ASAP1 | FHOD3 |  |
| MANF | KIF13B | RBFOX2 |  |
| FAU | CADM1 | WWOX |  |

